# *Cis-topic* modelling of single-cell epigenomes

**DOI:** 10.1101/370346

**Authors:** Carmen Bravo González-Blas, Liesbeth Minnoye, Dafni Papasokrati, Sara Aibar, Gert Hulselmans, Valerie Christiaens, Kristofer Davie, Jasper Wouters, Stein Aerts

**Author notes:** These authors contributed equally (Carmen Bravo González-Blas, Liesbeth Minnoye).

## Abstract

Single-cell epigenomics provides new opportunities to decipher genomic regulatory programs from heterogeneous samples and dynamic processes. We present a probabilistic framework called cisTopic, to simultaneously discover “cis-regulatory topics” and stable cell states from sparse single-cell epigenomics data. After benchmarking cisTopic on single-cell ATAC-seq data, single-cell DNA methylation data, and semi-simulated single-cell ChIP-seq data, we use cisTopic to predict regulatory programs in the human brain and validate these by aligning them with co-expression networks derived from single-cell RNA-seq data. Next, we performed a time-series single-cell ATAC-seq experiment after SOX10 perturbations in melanoma cultures, where cisTopic revealed dynamic regulatory topics driven by SOX10 and AP-1. Finally, machine learning and enhancer modelling approaches allowed to predict cell type specific SOX10 and SOX9 binding sites based on topic specific co-regulatory motifs. cisTopic is available as an R/Bioconductor package at http://github.com/aertslab/cistopic.

## Introduction

Genomic regulatory programs are driven by combinations of transcription factors that bind to cis-regulatory control regions, such as enhancers and promoters, thereby regulating the transcription of target genes. Unravelling the regulatory programs of different cell states can provide mechanistic insights into how these programs are encoded in the DNA sequence, how they are affected during disease, and how they can ultimately be exploited to manipulate cell fate, for example for cellular reprogramming. Although single-cell transcriptomics allows an unbiased detection of cellular diversity, reverse engineering the genomic regulatory code from the transcriptome remains a challenge. On the other hand, single-cell epigenomic techniques, such as single-cell ATAC-seq (scATAC-seq) (Buenrostro et al., 2015; Cusanovich et al., 2015), single-cell CUT&RUN (Hainer et al., 2018), or single-cell DNA methylome sequencing (Farlik et al., 2015), provide a more direct prediction of the genome-wide activity of enhancers and promoters, at single-cell resolution. These approaches, in particular single-cell chromatin accessibility profiling using scATAC-seq, allow the discovery of multiple cell types and regulatory states from a heterogeneous mixture of cells, such as a whole organism (Cusanovich et al., 2018), a whole organ (Lake et al., 2017), or an asynchronous dynamic process like differentiation (Corces et al., 2016; Pliner et al., 2017). These studies have provided extensive new insight into the diversity of chromatin landscapes within a tissue.

In comparison to single-cell transcriptomics, the computational analysis of scATAC-seq data is more challenging. This is mostly due to scalability and the higher sparsity of the data: a scATAC-seq dataset may harbour combinations from more than 100,000 potential regulatory sites –which results in extremely large matrices when profiling tens of thousands of cells–, but only a small subset of regions are detected as accessible in each individual cell (i.e. on average, only 10,000-20,000 deduplicated reads are obtained per cell (Table S1)). The current methods to analyse scATAC-seq data can be divided in two classes (Table S2). The first class consists of unsupervised methods such as scABC or Latent Semantic Indexing (LSI); in which, after representing the data in a lower dimensional space, cells with similar epigenomes are clustered (Cusanovich et al., 2015, 2018; Zamanighomi et al., 2018). Reads are then aggregated across all cells in a cluster to generate a pseudo-bulk profile, which is then used to identify differentially accessible regions between the clusters. A second class of methods consists of supervised methods that *a priori* aggregate all reads in a cell over pre-defined sets of genomic regions, called “*cistromes*” (e.g., ChIP-seq peaks of a transcription factor, or regions sharing a particular transcription factor motif or k-mer), such as chromVAR (Schep et al., 2017) and other, not yet peer-reviewed methods, such as BROCKMAN (de Boer and Regev, 2017) and SCRAT (Ji et al., 2017). Although this approach is effective to reduce the sparseness, it relies on pre-defined cistromes, which hinders the discovery of new regulatory programs. In addition, methods of both classes are optimised towards cell clustering, but do not provide a co-optimised grouping of regulatory regions.

Here, we develop cisTopic, an unsupervised Bayesian framework based on topic modelling, that allows simultaneous grouping of co-accessible regions into regulatory topics and clustering of cells based on their regulatory topic contributions. These “cis-regulatory topics” can be directly exploited for motif discovery to predict combinations of transcription factors, but also to explore dynamic changes in chromatin state. We benchmarked cisTopic using simulated data and concluded that this approach outperforms previously published methods in terms of accuracy, robustness and interpretability. We validated cisTopic by applying it to a previously published data set of 30,000 cells from the human brain (Lake et al., 2017), finding subpopulations in an unsupervised manner and in agreement with gene regulatory programs derived from single-cell transcriptomics data. In addition, we generate new scATAC-seq data and reveal dynamic changes in chromatin accessibility during melanoma phenotype switching *in vitro*, driven by the loss of SOX10. Finally, by comparing the SOX10 topics in melanoma with SOX9 and SOX10 topics in the brain, we propose a cooperative pioneering model for the SOXE (i.e. SOX8, SOX9 and SOX10) family members.

## Results

### Probabilistic topic modelling identifies cell states and reveals regulatory programs at single-cell resolution

We have developed cisTopic, a new method for the analysis of single-cell epigenomics data that allows the simultaneous identification of cell states and co-regulatory regions in an unsupervised manner (Fig. 1). The input for cisTopic is a binary accessibility matrix, with cells (i.e. objects) as columns and regulatory regions (i.e. features) as rows (in the case of single-cell methylation data, binary methylation scores) (Fig. 1a). Since this matrix is very sparse, we reasoned that Latent Dirichlet Allocation (LDA) (Blei et al., 2003), a robust Bayesian topic modelling method used to group objects addressing similar topics or themes, as well as grouping co-occurrent features into topics, could be applied to single-cell epigenomics data. Importantly, while existing methods rely on *hard clustering* (i.e., a feature or object will be uniquely assigned to one group), topic modelling assigns features to a group or topic with a certain probability, which means that the same feature can contribute to different groups, although with different strengths. In other words, compared to the discrete approach taken by clustering methods, the fuzzy clustering performed by topic models allows a feature (e.g. a regulatory region) to contribute to several groups or topics, and an object (e.g. a single cell) to be composed by different topics with different weights; resulting in less information loss.

**Figure 1.**
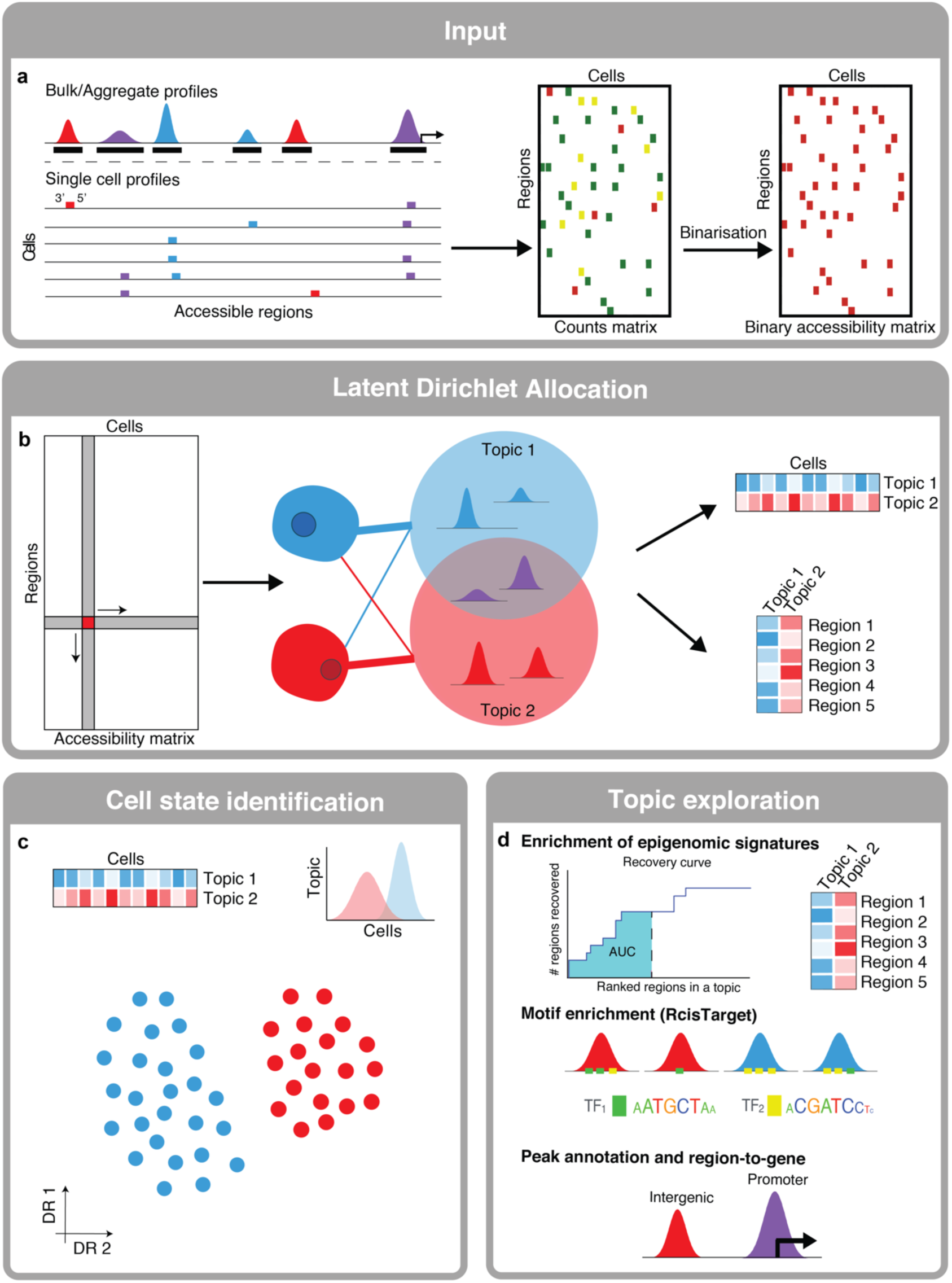
cisTopic workflow. **a.** The input for cisTopic is a binary accessibility matrix. This matrix can be formed from single-cell BAM files and a set of genome-wide regulatory regions (e.g., from peak calling on the bulk or aggregate data). **b.** Latent Dirichlet Allocation (LDA), using a collapsed Gibbs Sampler, is applied on the binary accessibility matrix to obtain the topic-cell distributions (contributions of each topic per cell) and the region-topic distributions (contributions of each region to a topic). Note that a region can contribute to more than one topic (represented by the purple peaks). **c.** The topic-cell distributions are used for dimensionality reduction (e.g. PCA, tSNE, diffusion maps) and clustering to identify cell states. **d.** The region-topic distributions can be used to predict the regulatory code underlying the topic. For example, topics can be compared with known epigenomic signatures using a recovery curve approach; regions can be annotated and linked to genes; and, after topic binarisation, enriched motifs can be identified via RcisTarget.

Importantly, LDA has a series of assumptions that are fulfilled in single-cell epigenomics data, such as non-ordered features (i.e. the order of regulatory regions is not relevant) and the allowance of overlapping topics (i.e. a regulatory region can be co-accessible with different other regions depending on the context; meaning that a region can participate in different regulatory programs depending on the cell type or state). In addition, compared to other topic modelling methods such as probabilistic LSI, LDA offers a probabilistic structure at the level of the objects by introducing Dirichlet priors over the topic contributions within the objects and does not lead to overfitting when increasing the size of the data set (Blei et al., 2003).

Several approaches have been proposed to estimate the probability distributions, such as maximising the probability of the features by estimating the feature-topic distributions using Expectation Maximization (which is slow and may converge to a local maxima) or Gibbs Sampling (Bishop, 2006; Blei et al., 2003). To maximise the performance of LDA, cisTopic uses a collapsed Gibbs Sampler (Griffiths and Steyvers, 2004), which allows to reduce the complexity of the model by only sampling the topic assignment of each feature per object without the need of sampling from the feature-topic and the topic-object distributions, reducing the exploration space. The probability of sampling from a specific topic is proportional to the contribution of that topic to the object and the contribution of that feature to the topic throughout the data set. These assignments are recorded through several iterations (after burn-in), and they can be used to estimate the feature-topic and the topic-object distributions.

Thus, we consider the accessible regulatory regions as features and cells as objects, and our aim is to simultaneously group regions that are co-accessible in topics and cluster cells based on the topic distribution of their accessible regions. By using LDA, two distributions are obtained, which correspond to (1) the probability of a region belonging to a cis-regulatory topic (region-topic distribution) and (2) the contributions of a topic within each cell (topic-cell distribution) (Fig. 1b). cisTopic includes functionalities for the biological interpretation of these distributions e.g. topic-cell distributions can be used to cluster cells and identify cell types (Fig. 1c); while region-topic distributions can be exploited to analyse the regulatory meaning of each topic (Fig. 1d). cisTopic is made available as a new R/Bioconductor package at http://github.com/aertslab/cistopic.

To benchmark cisTopic against the three published methods that are commonly used for scATAC-seq data analysis, namely Latent Semantic Indexing (LSI) (Cusanovich et al., 2015, 2018), chromVAR (Buenrostro et al., 2018; Johnson et al., 2018; Lareau et al., 2018; Liu et al., 2018; Mezger et al., 2018; Schep et al., 2017), and scABC (Zamanighomi et al., 2018); we simulated single-cell epigenomes from bulk H3K27Ac ChIP-seq profiles of 14 melanoma cell lines (Verfaillie et al., 2015). Eleven of these cell lines were published previously (GSE60666); while three additional profiles were generated in this study. To test the robustness of cisTopic towards sparsity, we ran several simulations varying the coverage per cell: from a range of 30,000-60,000 deduplicated reads per cell, down to 3,000-6,000 deduplicated reads per cell, similar to scATAC-seq coverage ranges found in literature (Table S1) (Fig. 2a; see *Methods*). We found that cisTopic is the most robust and accurate method to cluster cells (with an adjusted rand index (ARI) above 0.96, even at low read coverage), followed by scABC, LSI and chromVAR, respectively (Fig. 2b). Importantly, while previously existing methods only predict cell clusters, cisTopic simultaneously predicts regulatory regions that are important for each topic (i.e. other methods rely on *a posteriori* differential analysis of regions using the aggregated data per cluster (e.g. LSI and scABC, respectively) or start from *a priori* defined cistromes (e.g. chromVAR). On the melanoma H3K27Ac data, cisTopic reveals 2 general and 14 cell line specific topics (one for each cell line), as well as 2 topics that are shared across a subset of samples (Fig. 2c). One of these shared topics corresponds to the major melanoma cell line subtypes, namely the melanocyte-like, while the remaining corresponds to the mesenchymal-like subtypes (Hoek et al., 2006; Verfaillie et al., 2015). The genomic regions in these topics are enriched for AP-1 and TEAD motifs in the mesenchymal-like topic and SOX and E-box motifs in the melanocyte-like regions (Fig 2d), in agreement with earlier findings (Verfaillie et al., 2015). Furthermore, all the predicted topics from the simulated single-cell H3K27Ac ChIP-seq data can be confirmed by the corresponding bulk data (Fig. S1a,b). As expected, the general topics (accessible across all cell lines) are enriched for promoters; and the “low contribution topics” are formed mostly by “lowly accessible” regions and can be filtered out *a posteriori* (Fig S1c; Fig 2d). Next, we examined the capacity of all the tested methods to find rare subpopulations by reducing the number of cells for three of the cell lines by a 10-fold; and found that cisTopic also outperforms other methods in this aspect (Fig. S2). Finally, using previously published data sets of the hematopoietic system (Corces et al., 2016; Farlik et al., 2016), we confirmed that cisTopic also works on single-cell DNA methylation data and for trajectory analysis during differentiation (Fig S3, Fig S4).

**Figure 2.**
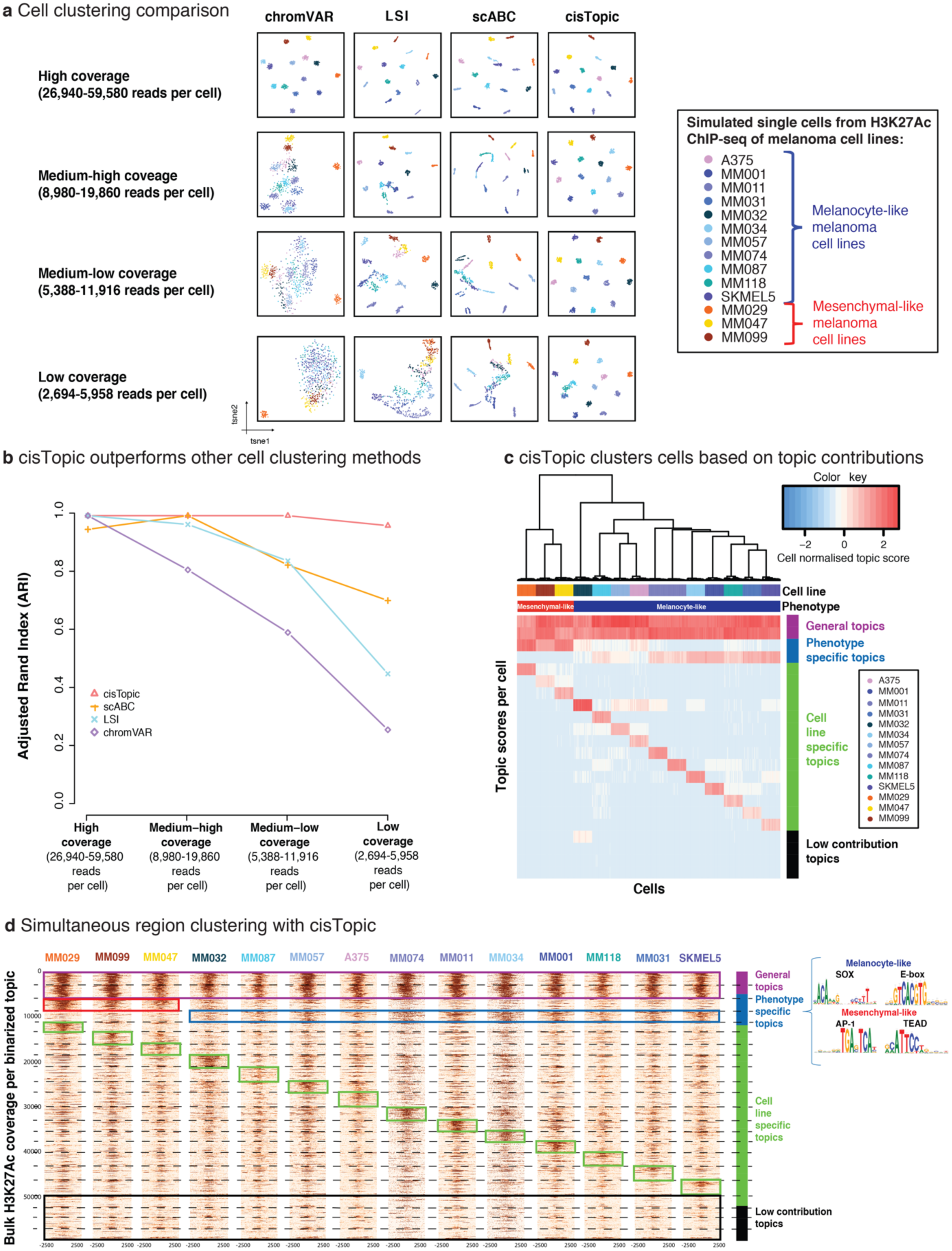
cisTopic outperforms other cell clustering methods, namely chromVAR, LSI and scABC; while simultaneously clustering regions into regulatory topics. **a.** Method comparison using semi-simulated single-cell H3K27Ac ChIP-seq data sampled from 14 bulk melanoma epigenomes with varying coverages. The tSNEs, coloured by cell line, were made using the cistrome enrichment matrix from chromVAR, the LSI matrix, the cell-to-landmark correlation matrix from scABC and the topic contributions per cell obtained with cisTopic. **b.** ARI for each method (chromVar, LSI, scABC and cisTopic) at each coverage, using the bulk epigenome of origin as ground truth and cluster assignments based on hierarchical clustering from the cistrome enrichment matrix (chromVAR), the LSI matrix, the cell-to-landmark correlation matrix (scABC) and the topic contributions per cell (cisTopic). cisTopic is the most robust method, even at low coverage. **c.** cisTopic clusters cells based on their topic contributions. Based on their distributions over the different cell populations, we found general, phenotype specific, cell line specific and low contributing topics. **d.** Coverage heatmaps of bulk H3K27Ac data to validate the predicted regions per topic (see *Methods)*. Each binarised topic is represented between the dashed horizontal lines, and within each topic, the regions are ordered by descending topic score. Topic regions show the expected patterns in the bulk data (expected patterns are surrounded by squares). Key motifs found enriched in the phenotype specific regions by RcisTarget are shown (right).

In conclusion, cisTopic not only defines cell states more accurately than existing methods, but also discovers meaningful regulatory topics that yield insight into cell-type specific regulatory programs.

### cisTopic identifies robust cell types and gene regulatory networks in the human brain

Next, we applied cisTopic to a large and biologically complex scTHS-seq data set (obtained by single-cell Tn5 Hypersensitivity Sequencing, similar to scATAC-seq) with 34,520 single cells from the human brain (Lake et al., 2017). This data set contains cells from the cerebellum, frontal cortex and visual cortex from three patients; with a total of 287,381 accessible regulatory regions. Based on the log-likelihood in the last iteration of the models, we selected the optimal number of regulatory topics to be 23 (see *Methods*; Fig. S5). Using the topic-cell distributions, we were able to cluster the cells according to the major brain cell types: excitatory neurons (Ex), inhibitory neurons (In), cerebellar granule (Gran) cells, endothelial cells (End), astrocytes (Ast), oligodendrocytes (Oli), oligodendrocyte precursor cells (OPCs) and microglia (Mic) (Fig. 3a; Fig. S6a-e). After selecting the representative regions per topic by fitting a gamma distribution on the region-topic distributions (see *Methods*), we used RcisTarget (Aibar et al., 2017) to predict enriched motifs in each topic. For example, SOX and NFIA/B motifs are enriched in enhancers that are specifically accessible in astrocytes; SOX and OLIG motifs in the oligodendrocyte regulatory topic and NEUROD in the granule cell specific topic (Fig. S7). SOX9 and NFIA are known as key transcription factors during astrocyte development and maintenance, and their combined over-expression is sufficient to trans-differentiate fibroblasts into astrocytes (Caiazzo et al., 2015; Kang et al., 2012; Sun et al., 2017; Wilczynska et al., 2009). SOX10, OLIG1 and OLIG2 are master regulators of oligodendrocyte development (Wegner and Stolt, 2005; Yu et al., 2013; Zhou and Anderson, 2002). and NEUROD1/2 is a marker of granule cell differentiation (Miyata et al., 1999). In fact, several regions in the vicinity of the *NEUROD1* gene are highly accessible in the cerebellum, where granule cells reside, as compared to the visual and the frontal cortex (Fig. S8). Finally, the predicted regulatory topics could be further validated by GO enrichment using GREAT (McLean et al., 2010)), finding “myelination” (GO:0042552, p-value: 10^-23^) for the oligodendrocytes topic, “glial cell fate commitment” (GO:0010001, p-value: 10^-7^) for the astrocytes, and "regulation of sensory perception of pain" (GO:0051930, p-value: 10^-8^) for the granule cells. Indeed, cerebellar granule cells are involved in sensory cognition (Bing et al., 2015).

**Figure 3.**
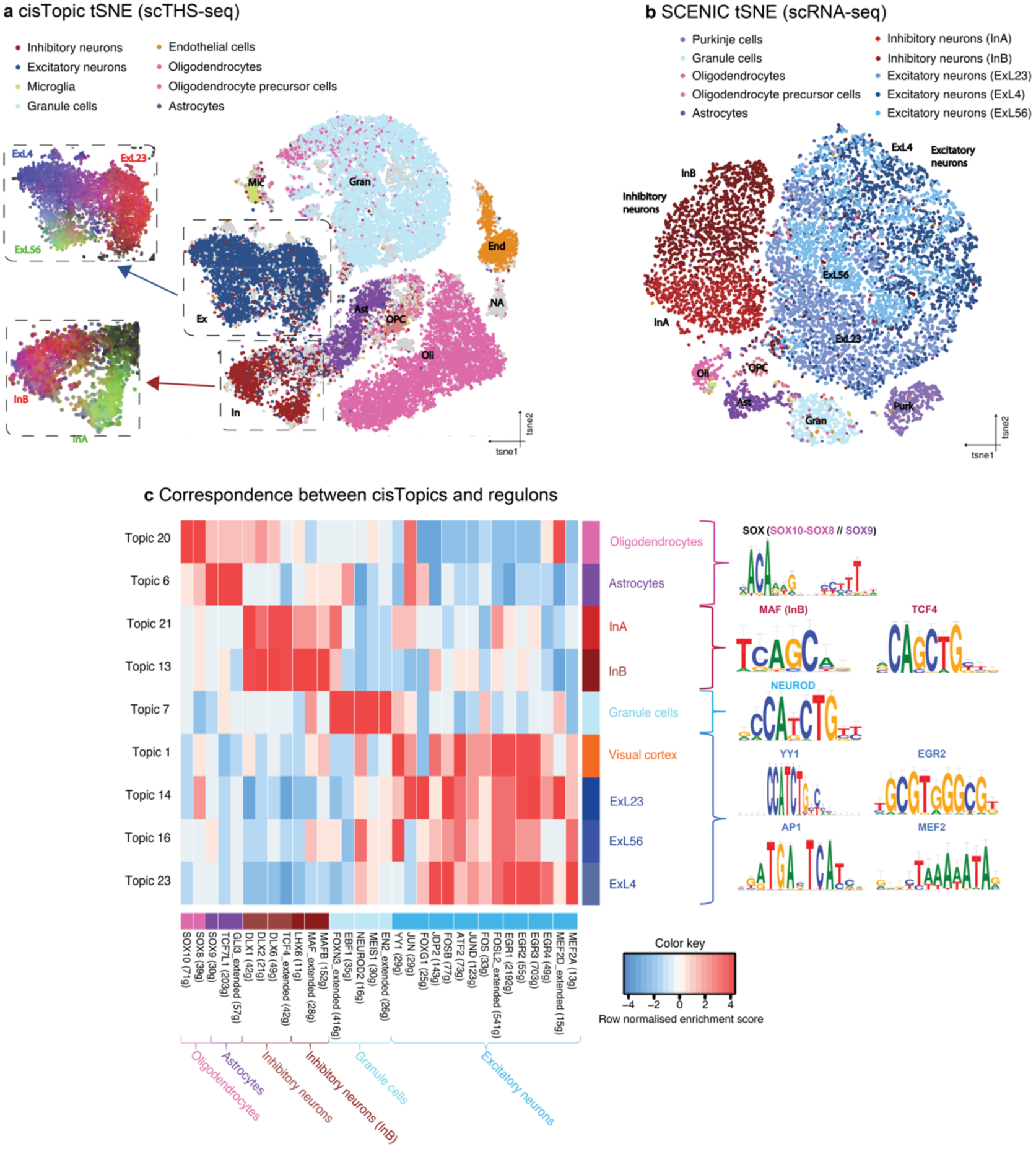
cisTopic reveals major cell types and subpopulations in the human brain and summarises regulatory programs underlying the transcriptome. **a.** cisTopic tSNE based on topic-cell contributions from the analysis of the scTHS-seq data. cisTopic identifies the main cell types but also subpopulations in interneurons (InA and InB) and excitatory neurons (ExL23, ExL4 and ExL56). Contributions of the subpopulation specific topics are represented by RGB encoding. **b.** tSNE based on regulon enrichment obtained using SCENIC (Aibar et al., 2017) on the scRNA-seq data from the same tissue. **c.** Correspondence between cisTopic topics and SCENIC regulons. The motifs shown are found in both the matching regulon and the topic.

Interestingly, cisTopic also revealed heterogeneity within the interneurons, with two distinct subtypes (InA and InB), from which one (InB) is enriched for MAF motifs (Fig. S7). In the MAF-enriched topic, targets such as ARX, LHX6, SOX6 and DLX1 are found; which, together with MAF and MAFB are markers for the Medial Ganglionic Eminence derived interneurons (Chen et al., 2017; Lake et al., 2016). In the second interneuron topic (InA population) PCP4, ISL1, SP8 and VIP are found, which are markers for the Caudal/Lateral Ganglionic Eminence derived interneurons (Chen et al., 2017; Lake et al., 2016). Also, based on the cisTopic analysis, three subtypes of excitatory neurons can be distinguished (Fig. 3a, Fig. S7). These represent different neuronal layer positions within the cortex, namely layers II and III (ExL23), IV (ExL4) and V and VI (ExL56). These subpopulations had been already reported by Lake *et al.* based on scRNA-seq data (2017); however, cisTopic is able to distinguish them directly from scATAC-seq data, without the need of other data (Fig. S6f,g).

To further validate the predicted regulatory topics, we explored the relationship between cell type specific regulatory regions and cell type specific gene expression. To do so, we used the matching scRNA-seq data generated by Lake et al. (2017) on the same human brain tissues (15,884 cells). We run SCENIC to infer gene regulatory networks and cluster cell types from this data, predicting 250 “regulons”, whereby each regulon consists of a transcription factor and its predicted target genes based on co-expression and motif enrichment (Aibar et al., 2017). Cell clustering using these 250 regulons identified the major cell types of interneurons and excitatory subpopulations at the same resolution as cisTopic (Fig. 3b). By comparing the cell type specific scRNA-seq regulons with the scATAC-seq topics (see *Methods*), we found a strong agreement for a range of transcription factors. For example, SOX8 and SOX10 regulons match the oligodendrocyte topic; and the SOX9 and GLI3 regulons that correspond with the astrocyte topic. Likewise, the DLX regulons match with the interneurons topics; and specifically, LHX6, MAF and MAFB regulons correspond with InB interneuron topic; NEUROD2 regulons match with cerebellar topics; and AP-1, EGR and MEF2 regulons with excitatory neuron topics (Fig. 3c; Fig. S9). Importantly, the predicted transcription factors controlling the cis-regulatory topics (based on motif enrichment) and the corresponding expression-based regulons show strong agreement with literature (Chen et al., 2017; Flavell et al., 2008; Gashler and Sukhatme, 1995; Kaczmarek, 2002; Lake et al., 2016; Miyata et al., 1999; O’Donovan and Baraban, 1999; Petrova et al., 2013; Sun et al., 2017; Turnescu et al., 2018).

In conclusion, cisTopic reveals with high sensitivity cell states in large and heterogeneous data sets such as the human brain. Furthermore, the defined regulatory topics represent biologically relevant gene regulatory networks as demonstrated by the enrichment of motifs related TFs important in the defined cell types and correspondence between single-cell epigenomes and single-cell transcriptomes.

### cisTopic maps a dynamic regulatory landscape downstream of SOX10 in melanoma

Next, we applied cisTopic to investigate dynamic changes in chromatin accessibility during a cell state transition in melanoma cells. *In vitro* studies of melanoma lines (Bittner et al., 2000; Hoek et al., 2006; Restivo et al., 2017), and later *in vivo* studies (Eichhoff et al., 2010; Hoek et al., 2008; Wouters et al., 2014), have identified two stable subpopulations in melanoma, characterised by very distinct transcriptomes: a ‘melanocyte-like’ state, with high expression of the melanocyte lineage specific transcription factor MITF (Hoek et al., 2006) as well as high SOX10 and PAX3 (Scholl et al., 2001; Shakhova et al., 2012); and an ‘invasive’, drug-resistant, mesenchymal-like state with low levels of MITF, high levels of genes involved in TGFb signalling and governed by AP-1 and TEAD transcription factors (Hoek et al., 2008; Verfaillie et al., 2015). The transcription factor SOX10, a major regulator of neural crest development and melanocytic differentiation (Harris et al., 2011; Kellerer, 2006), plays an important role in maintaining the melanocyte-like state, as loss of SOX10 has previously been shown to upregulate invasive genes such as *JUN*, *AXL*, and *SOX9* (Shaffer et al., 2017; Shakhova et al., 2012; Verfaillie et al., 2015), increase vemurafenib resistance (Sun et al., 2014), and induce a stable resistant state regulated by AP-1 (Shaffer et al., 2017). To study the regulatory dynamics of the switch from the melanocyte-like state towards the mesenchymal-like state, we performed a time series experiment after knockdown (KD) of SOX10 in two melanocyte-like melanoma cultures (MM057 and MM087) (Gembarska et al., 2012; Verfaillie et al., 2015) (Fig. 4a). As it is currently unknown whether melanocyte-like cells within one population follow the same regulatory path during the phenotype switch, we performed scATAC-seq, using the Fluidigm C1, at 0, 24, 48 and 72 hours after SOX10 KD (Fig. 4a). After filtering out cells with low signal TSS-aggregation plots, we obtained 598 cells in total with an average of 54,343 reads per single cell, and a total of 78,262 peaks over all conditions (see *Methods*). We also performed bulk OmniATAC-seq (Corces et al., 2017) on the same time points and cell lines to validate the quality of the scATAC-seq data. Aggregated profiles of scATAC-seq data closely resemble bulk OmniATAC-seq data in the same conditions (Fig. S10a,b) and there was a clear correlation between the corresponding conditions in bulk and single-cell ATAC-seq samples (average correlation coefficient of 0.83, Fig. S10c). The effectiveness of the transcriptional switch was confirmed by the loss of accessibility over time at promotors and enhancers near marker genes of the melanocyte-like state, such as *DCT* and *TYR*, genes involved in melanin production (Bernd et al., 1994; Iozumi et al., 1993), and *ERBB3* (Buac et al., 2011) (Fig. 4b and Fig. S11a); and by gain of accessibility of mesenchymal-like regions such as a *CLDN4* enhancer (Fig. S11b).

**Figure 4.**
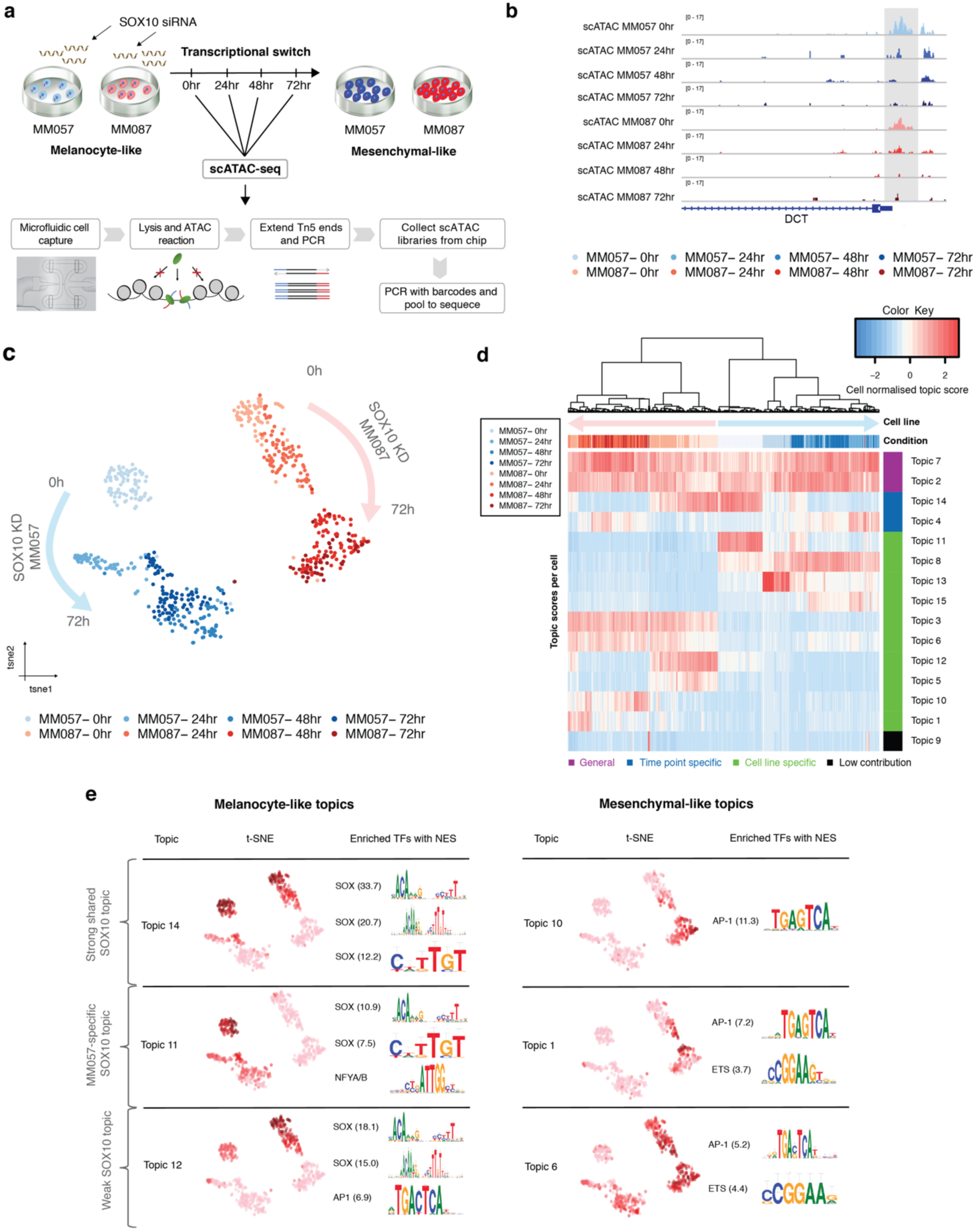
scATAC-seq during an EMT-like transition triggered by SOX10 knockdown in melanoma. **a.** scATAC-seq was performed with the Fluidigm C1 on two melanoma lines (MM057 and MM087) during a SOX10-KD-induced transcriptional switch from a melanocyte-like to a mesenchymal-like state at four time points (0, 24, 48 and 72 hours post-SOX10-KD). **b.** Profiles of scATAC-seq aggregates per condition in the region surrounding *DCT*, a SOX10 target gene that loses accessibility at the SOX10 binding site during the transition. **c.** tSNE-representation (598 single cells) generated by cisTopic using the cell-topic distributions showing the dynamics of the switch in MM057 (blue) and MM087 (red) at the four different time points after SOX10-KD. **d.** cisTopic heatmap of topic distributions within the single cells. Several classes of topics are identified, namely general topics, time point and cell line specific topics. **e.** cisTopic identifies several melanocyte-like and mesenchymal-like regulatory topics, represented here as t-SNEs coloured by the topic score together with representative enriched TF motifs per topic (ordered by Normalised Enrichment Score (NES)).

When we applied cisTopic to this dataset we found that a model with 15 regulatory topics best represented the data (Fig. S12a; Fig. 4c,d). A few topics represent genomic regions that are ubiquitously accessible, across both cell lines, and across all time points (topic 2 and 7) (Fig. 4d). Regions with high probability of belonging to these topics are strongly enriched for promoters (Fig. S10b) and for SP1 and NFY motifs, two common promoter motifs (Fig. S13) (Dynan and Tjian, 1983; Li et al., 1992). The remaining topics are mostly specific for a cell line, specific for a time point, or specific for a particular combination of cell line and time point (Fig. 4d; Fig. S13). Several topics represent regions that become accessible at later time-points after SOX10 knockdown (e.g. topic 10, 1 and 6) (Fig. 4d,e; Fig. S13; Fig. S14). Particularly, topic 10 is reminiscent of the previously described invasive/mesenchymal-like epigenome (Verfaillie et al., 2015) (Fig. S15) and genes near topic 10 regions are involved in cell migration, e.g., *EGFR*, *TGFB2*, *TGFBR2* and *AXL*. Motif discovery on the regions composing topic 10 identified motifs linked to the AP-1 transcription factor family, such as JUNB, JUND and FOS (Fig. 4e), as well as an enrichment of ChIP-seq peaks for TEADs (Fig. S15). As AP-1 and TEAD are known regulators of melanoma cells in the mesenchymal-like state (Shaffer et al., 2017; Verfaillie et al., 2015), these results agree with previous findings. We note that all cells undergo similar epigenomic changes during the transition (Fig. 4c), indicating that, with the resolution obtained by this experiment, there is no heterogeneity in the way the chromatin changes during the transition.

cisTopic also predicts three topics that show a decline in accessibility during the state transition. The strongest of these topics, topic 14, is shared between the two tested cell lines (Fig. 4d,e; Fig. S14). Two additional declining topics are specific to either MM057 (topic 11) or MM087 (topic 12) (Fig. 4d,e; Fig. S14-S16). Motif discovery revealed that the enhancers composing these three ‘melanocyte-like’ topics were highly enriched in motifs linked to the SOX transcription factor family (Fig. 4e). Given that we knockdowned SOX10 and its role in the melanocyte-like state, SOX10 is the most likely candidate transcription factor to bind these regions in the melanocyte-like state. Indeed, by comparing these topics with previously published SOX10 ChIP-seq data obtained from a melanocyte-like melanoma line (Laurette et al., 2015), we observed strong SOX10 ChIP-seq signal on the regions belonging to these topics (Fig. S16a). For example, several known, experimentally validated SOX10 target enhancers, such as binding sites near *ERBB3* (Prasad et al., 2011), *MIA* (Graf et al., 2014), *TYR* (Murisier et al., 2007) and *DCT* (Potterf et al., 2001) all contain a topic 14 region overlapping with a SOX10 ChIP-seq peak (Fig. S11a; Fig. S16c). Importantly, the finding that SOX10 KD results in chromatin closing of SOX10 enhancers (topic 11, 12 and 14) suggests that SOX10 is a chromatin modifier. In agreement with this, loss of SOX10 directly impacts chromatin accessibility (i.e. regions decreasing in accessibility are directly linked to SOX based on motif enrichment); and higher SOX10 protein levels in MM087 compared to MM057 result in longer residence times at shared SOX10 targets, with increased accessibility of peaks in MM087 as well as a slower SOX10-KD-induced state transition (Fig. S14; Fig. S16d; Fig. 4c).

This study shows that scATAC-seq data during state transitions can be used together with cisTopic to uncover the regulatory dynamics of biological processes, such as the EMT-like transition in melanoma induced by knockdown of the transcription factor SOX10. Our results show that all cells follow a common path during this switch, which involves ~1000 functional SOX10 enhancers that decline in accessibility during the transition, showing that SOX10 has an effect on the chromatin landscape.

### A cooperative-pioneer enhancer model for SOXE transcription factors

Regulatory topics identified by cisTopic represent high-quality sets of functional enhancers that allow in-depth analysis of the composition of transcription factor binding sites. Indeed, the accuracy of SOX10 enhancer prediction from cisTopic is comparable to the accuracy of ChIP-seq, since the enrichment of SOX motifs within the SOX10 topic regions is comparable to the enrichment of SOX motifs in SOX10 ChIP-seq data (NES score of 33.74 for the shared SOX10 topic in melanoma, compared to 33.79 for SOX10 ChIP-seq in melanoma). We reasoned that cis-regulatory topics can be used to decipher transcription factor specific enhancer architectures. Particularly, we compared three different SOXE topics (which comprise SOX8, SOX9, and SOX10 (Wright et al., 1993)); namely the oligodendrocyte (SOX10) and the astrocyte topic (SOX9) from the human brain data set (Fig. S6, S7) and the shared SOX10 topic during the melanoma EMT-like transition (topic 14, Fig. 4e). In these three topics, the top enriched motif is the same SOX dimer (with NES scores of 20.00, 8.78 and 33.74 for oligodendrocytes, astrocytes and melanoma, respectively). For each of the three topics, we selected the subset of regulatory regions enriched for SOX motifs (see *Methods*). These three sets of regions are largely unique (~17% overlap on average) (Fig. 5a). The distinct use of SOX10 enhancers between cell types is confirmed by plotting the melanoma scATAC-seq signal on the oligodendrocyte and astrocyte SOX cistromes, as only a limited subset of the brain targets is accessible in melanoma (Fig. 5b). The finding that SOXE factors regulate distinct targets in different cell types is expected, since they play different roles depending on the cell type (Harris et al., 2011; Kellerer, 2006; Stolt et al., 2002). Genes linked to SOX regions exclusively found in the melanoma SOX topic are significantly enriched for the GO term “pigmentation” (GO:0043473, p-value: 10^-3^); genes linked to SOX regions exclusively found in the oligodendrocyte SOX topic are significantly enriched for the GO term “myelination” (GO:0042552, p-value: 10^-14^); while genes linked to SOX regions exclusively found in the astrocyte SOX topic are enriched for the GO term “gliogenesis” (GO:0042063, p-value: 10^-8^). Note that the entire set of melanoma regions is also accessible in melanocytes, as shown by DNAseI-seq data in melanocytes (Fig. S17), suggesting that there are no “ectopic” functional SOX10 binding sites in melanoma beyond those that exist in melanocytes. Therefore, we can exploit the SOX10 topic in melanoma to investigate the SOX enhancer architecture in melanocytes.

**Figure 5.**
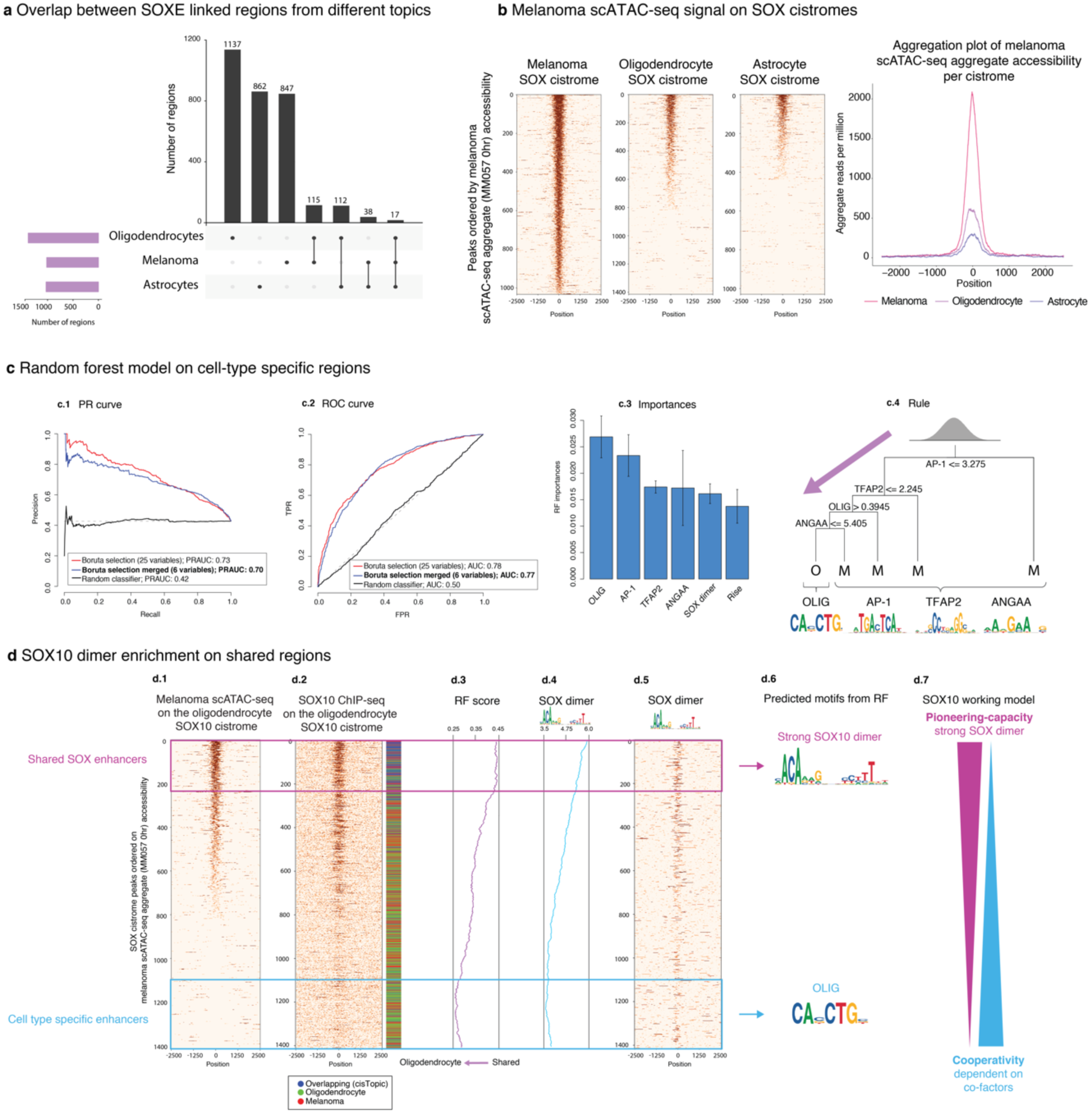
Comparison of SOXE cis-topics between cell types. **a.** Number of unique and overlapping regions between the oligodendrocyte, melanoma and astrocyte SOX cistromes. **b**. Heatmaps and aggregation plot showing melanoma scATAC-seq signal on the melanoma, oligodendrocyte and astrocyte SOX cistrome regions. Cistrome peaks are ranked according to their scATAC-seq accessibility in melanoma (MM057, 0 hours after SOX10-KD). **c**. Random forest model to discriminate between melanoma and oligodendrocyte specific SOX regions. **c.1**. Precision Recall (PR) and **c.2.** Receiver Operating Characteristic (ROC) curves for different Random Forest models using either 25 variables after Boruta selection, 6 variables after merging correlating variables from Boruta, or a random classifier. **c.3**. Variable importances for the RF model with merged motif features **c.4**. Representative rule extracted from the Random Forest model with InTrees (Deng, 2014). Each root represents a decision point with a rule based on one of the variables (CRM scores) used in the RF model (if the rule is fulfilled, the left path is taken). The leaves represent the class assigned (O: Oligodendrocytes; M: Melanoma). The motifs used in the RF model with 6 variables are shown under the rule tree, showing whether they are either melanoma or oligodendrocyte specific. **d. d.1**. Heatmap showing scATAC-seq signal and **d.2**. SOX10 ChIP-seq on the oligodendrocyte SOX cistrome region ranked according to their scATAC-seq accessibility in melanoma. The colour bar next to the heatmaps represents whether the regions were either overlapping with regions with the other SOX cistromes (blue), whether they were correctly classified as an oligodendrocyte region using the rule extracted with Intrees (green) or whether they were misclassified as melanoma regions (red). Regions that are not unique to the oligodendrocyte SOX10 cistrome (blue) are enriched on top of the heatmap, meaning that they are also accessible in melanoma, and have higher SOX10 ChIP-seq signal. These regions are highlighted by the pink box as shared SOX enhancers. Regions that are specific to oligodendrocytes are enriched at the bottom of the heatmaps and are highlighted by the blue box. **d.3.** RF scores for the heatmap regions. **d.4.** SOX dimer CRM scores for the heatmap regions. **d.5.** Heatmap representing SOX10 CRMs in the sequences. **d.6.** Logos of motifs enriched in the shared SOX enhancer and the oligodendrocyte-specific enhancers as found by RF. **d.7.** Representation of the potential model. Shared regions are enriched for SOX dimers, while cell type-specific regions are enriched for co-factors.

**Figure 6.**
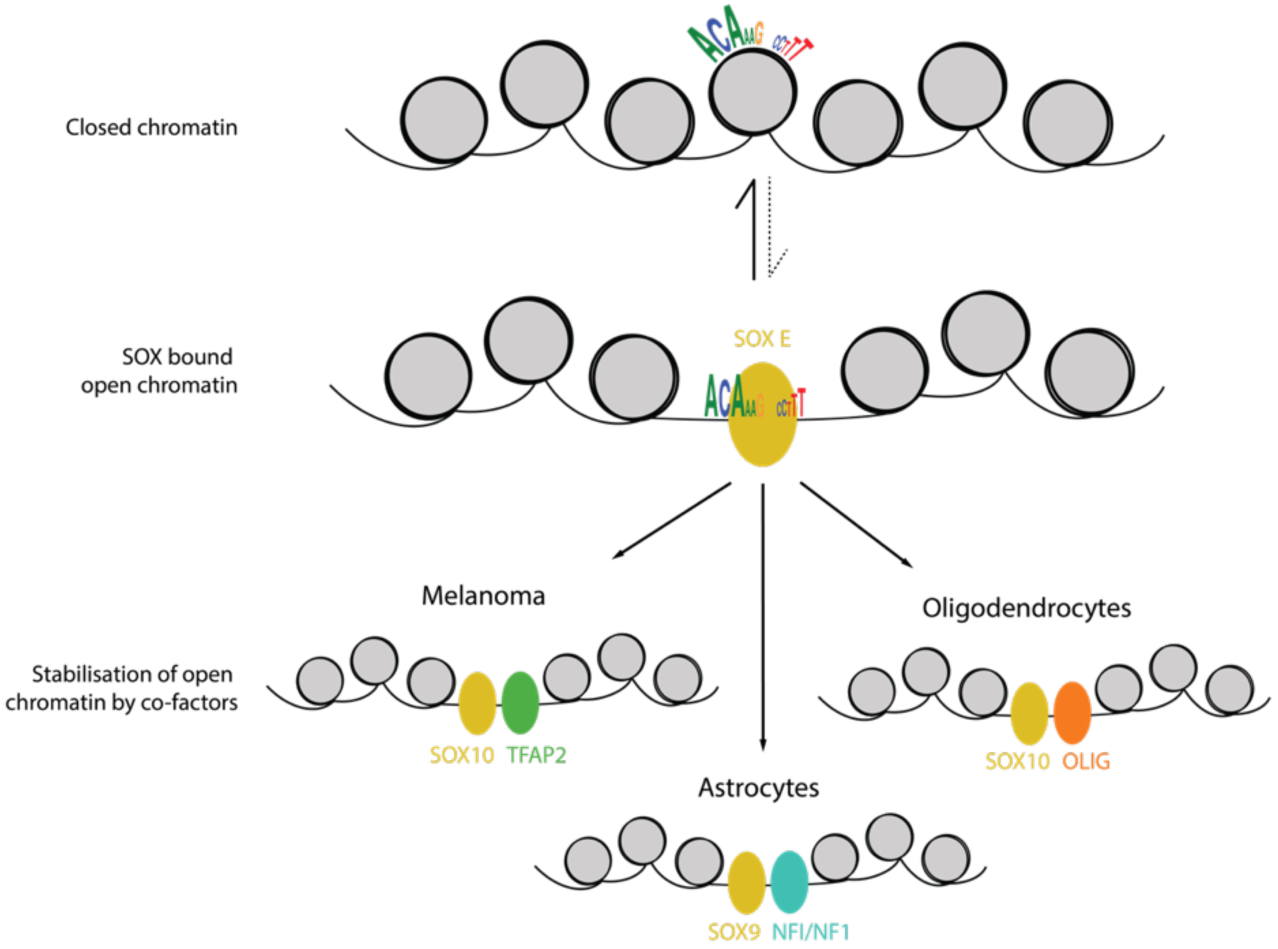
Quantitative pioneering function of SOXE proteins depends on binding stabilisation by cell-type specific co-factors. SOXE proteins are able to recognise their binding sites; however, when the binding site is not strong enough, they require the help of additional cell type-specific co-factors, such as TFAP2 in melanoma, OLIG in oligodendrocytes and NFI in astrocytes, to be stable.

How the specificity of these regulatory programs is achieved is largely unknown, although previous studies investigating SOXE enhancer codes have suggested that cooperativity with other TFs is common (Hou et al., 2017; Kondoh and Kamachi, 2010; Wilson and Koopman, 2002). To identify the sequence features that result in SOXE cell type specific programs, we compared the non-overlapping regions between the selected SOXE cistromes in a pairwise manner, using a Random Forest model (see *Methods*). Here, we focus on the comparison between the two SOX10 cistromes (in melanoma and oligodendrocytes), while comparisons between SOX10 and SOX9 cistromes are shown in Fig S21. As candidate features we used known and *de novo* motifs (from the cisTarget motif collection (Herrmann et al., 2012; Imrichová et al., 2015) and Homer (Heinz et al., 2010) and RSAT *peak-motifs* (Thomas-Chollier et al., 2011, 2012), respectively) and DNA shape measurements from GBshape and Kaplan et al. (Chiu et al., 2015; Kaplan et al., 2009) (Fig. S18). The motifs were scored in the regions using Cluster-Buster (Frith et al., 2003), selecting as features the best Cis-Regulatory Module (CRM) score per region; while for DNA shape measurements we used the average value in ±250bp from the centre of the regions (Fig. S18). A likelihood ratio test between the groups, resulted in 3,816 features selected (FDR adjusted p-value < 0.05). These features were used as input for Boruta (Kursa and Rudnicki, 2010), which found 25 informative features (see *Methods*). Among these 25 features there were several similar and correlated motifs (e.g. multiple E-box PWMs), which we merged into one Hidden Markov Model score using Cluster-Buster (Frith et al., 2003), resulting in a final model containing 6 features. The performance of the RF model with only these 6 features achieved a similar performance compared to the model using all 25 Boruta features, with an Area Under the Precision Recall (AUPR) of 0.70 and an Area Under the Curve (AUC) of 0.77 (Fig. 5c1,c2). This simplified model suggests that melanoma-specific SOX10 binding is determined mainly by co-binding of factors from the TFAP2 (AP-2) and AP-1 family; while oligodendrocyte-specific binding is determined by co-binding of bHLH family members of the CA<u>GC</u>TG type, likely reflecting OLIG (Fig. 5c3,c4) (Mazzoni et al., 2011; Yu et al., 2013). In addition, a *de novo* motif, AnGAA, is found enriched within the SOX melanoma cistrome. TFAP2 is a plausible candidate for a co-regulatory factor of SOX10 in melanoma and melanocytes given its important role, along with SOX10, in controlling melanocyte fate (Seberg et al., 2017). Indeed, the enhancers with predicted SOX10 and TFAP2 binding sites show strong overlap with previously published TFAP2A ChIP-seq peaks in melanocytes (Fig. S19) (Seberg et al., 2017). Although the MITF motif was not selected as top feature, the melanoma SOX enhancers also show strong overlap with MITF-bound regions found by ChIP-seq in melanoma (Fig. S19) (Laurette et al., 2015). Interestingly, co-occurrence of TFAP2 and MITF binding sites has been previously reported (Seberg et al., 2017). The AP-1 motif is also very strongly enriched in the melanoma-specific enhancers, and members of the AP-1 family, like JUN and FOS, are expressed in melanoma and melanocytes, while markedly absent from oligodendrocytes (Fig. S20). Finally, we find a predicted DNA shape feature enriched in the melanoma SOX enhancers, namely *rise*, which is positively correlated with other sequence features such as GC content, nucleosome occupancy, hydroxyl radical cleavage and propeller twist; and negatively correlated with helix twist (Fig. S21). This may suggest that cell type specific SOX10 binding may, next to distinct co-factors, also require a specific sequence environment. A similar analysis of SOX10 enhancers versus SOX9 enhancers (oligodendrocyte versus astrocyte and melanoma versus astrocyte cistromes, respectively) revealed features such as NFIA/B motifs strongly enriched in astrocyte enhancers, which is in agreement with literature (Kang et al., 2012) (Fig. S22).

Next, we investigated the SOX regulatory regions that show shared accessibility across multiple cell types. As expected, these shared regions cannot be classified into one of the cell types with our trained RF model, having a RF score of around 0.5 (Fig. 5d3). When comparing these shared regions to cell type specific regions, we found that shared enhancers show higher SOX10 ChIP-seq signal (Fig 5d1,2), and stronger SOX10 dimer motifs (LRT FDR p-value: 10^-9^) compared to the cell type specific enhancers (Fig 5d4,5,6). Interestingly, monomer motifs are not enriched in the shared regions (LRT FDR p-value: 0.78). Altogether, these findings indicate that shared enhancers could be bound by SOX10 homodimers alone (or for example SOX10-SOX8 heterodimers), with longer residence time; whereas cell type specific enhancers have weaker SOX10 dimer motifs, avoiding activation in the wrong cell type, but more prevalent co-regulatory motifs to regulate their distinct function (Fig. 5d7).

We further tested this hypothesis using enhancer-reporter assays (Fig. S23). The DCT enhancer, which is specific for melanoma, has strong predicted TFAP2, AP-1, and MITF CRM scores (SOX dimer: 4.50; TFAP2: 2.43; AP-1: 0.269; MITF: 5.77). Mutating the MITF binding sites abolishes the DCT enhancer activity. On the other hand, the EDNRB enhancer, which is also accessible in the brain, has stronger SOX10 binding sites (SOX dimer: 8.56), but weaker co-factor CRMs (AP-1: 0; TFAP2: 1.44; MITF: 0.905). Indeed, mutating E-boxes in the EDNRB enhancer did not have a significant effect on enhancer activity (Fig. S23). This indicates that cell type specific enhancers, with strong co-factor motifs, are more prone to losing their activity when the specific co-factor is not present; whereas enhancers that are accessible in several cell types and contain strong SOX10 dimer motifs are nearly unaffected by loss of the co-factor. Note that for both enhancers, mutating the SOX10 motif completely abolishes enhancer activity (Fig. S23).

In conclusion, regulatory topics identified by cisTopic, based only on single-cell ATAC-seq data, represent functional enhancers of high quality that can be used to decipher the regulatory logic of enhancer specificity. When applied to study SOXE enhancers, we found that certain SOX regulatory regions are specifically accessible in a given cell type; while others are accessible across systems. Using Random Forest models we could distinguish between melanoma-and oligodendrocyte-specific SOX enhancers based on motifs of cooperatively bound transcription factors; while we found that shared SOX10 enhancers show a preference for SOX10 dimer motifs and may be driven by pioneering activity of SOX10.

### Discussion

Single-cell epigenomics, particularly single-cell ATAC-seq, yield unprecedented insight into chromatin landscapes of individual cells. However, for each individual cell only a very limited number of accessible regions can be sampled, i.e. ~10% of all open regions. In other words, the data obtained from a single cell cannot be used directly to predict which genomic regions are accessible. To overcome this problem, currently available methods either aggregate ATAC-seq reads across a set of “similar” cells, following a cell clustering based on dimensionality reduction (e.g., scABC and LSI (Cusanovich et al., 2015, 2018; Zamanighomi et al., 2018)); or alternatively, aggregate ATAC-seq reads per cell across a predefined set of genomic regions (de Boer and Regev, 2017; Ji et al., 2017; Schep et al., 2017). Although these solutions have been shown to be satisfactory to cluster cells and to identify cell types, they do not allow the *ab initio* identification of co-regulatory regions (or *cis-regulatory topics*). Here, we have shown that Bayesian topic modelling, particularly LDA, allows the simultaneous discovery of *cis-topics* and cell types. LDA groups features into topics with a certain score (i.e. a feature can belong to several topics with different preferences); and objects can be represented as a mixture of topics. Compared with the discrete approach taken by conventional clustering methods (i.e. a feature or object can only belong to one group), this algorithm results in less information loss.

Topic modelling has been previously used in other fields for dealing with noisy data, such as text mining, image processing and forensics (Blei et al., 2003; Kuang et al., 2017; Rasiwasia and Vasconcelos, 2013). We have extrapolated this framework to single-cell epigenomics, by considering cells as objects; genomic regions as features; and cis-regulatory topics (or cis-topics) as topics. In agreement with the high accuracy of LDA in other fields, cisTopic groups cells into cell types and cell states, even when data is extremely sparse, with higher accuracy than currently published methods; and simultaneously group genomic regions into cis-topics; something that, to our knowledge, has not been shown before. Furthermore, cisTopic also includes functionalities to explore the output of LDA for biological interpretation. For example, topic contributions within cells can be used for cell type identification (i.e. clustering, tSNE), while regulatory topics can be used to decipher cell-state specific regulation (i.e. motif enrichment and machine learning).

The performance of cisTopic was confirmed on simulated H3K27Ac ChIP-seq data, which we believe represents a relevant test case, given that the recently developed single-cell CUT&RUN (an alternative to ChIP-seq to profile TF binding or histone modifications in single cells) will likely be widely adopted (Hainer et al., 2018). Our results on 14 melanoma cell lines showed that cell clustering is 96% accurate even with as few as 3,000 reads per cell. More importantly, the predicted topics reveal meaningful regulatory programs, some cell-type specific, and some shared by cell lines from the same melanoma subtype.

Single-cell epigenomics data sets are becoming increasingly large: recent data sets obtained from the Drosophila embryo (Cusanovich et al., 2018) and the human brain (Lake et al., 2017) contain more than 30,000 cells. Thanks to a binarisation step, and the use of a collapsed Gibbs sampler, cisTopic is computationally efficient to analyse such large data sets. By increasing the number of cells, but also by combining multiple single-cell omics layers, like scATAC-seq and scRNA-seq, the power to detect new and rare cell types and subpopulations from heterogeneous tissues becomes more and more feasible. Previously, Lake et al. (2017) combined the analysis of scATAC-seq and scRNA-seq data using Gradient Boosting Machines, which allowed them to identify subpopulations of inhibitory and excitatory neurons at the chromatin level. Interestingly, in our study, using cisTopic, we could identify the same subpopulations *ab initio* from uniquely the scATAC-seq data. The predicted topics and candidate transcription factors were then confirmed *a posteriori*, through an independent network analysis of the corresponding scRNA-seq data. The finding that epigenome-based cis-topics correspond to gene regulatory networks is encouraging for future studies, particularly when single-cell multi-omics strategies can be up-scaled (Angermueller et al., 2016; Hu et al., 2016; Pott, 2016).

scATAC-seq has been mainly applied to complex tissue samples, such as the hematopoietic system, the human and mouse brain, and the Drosophila embryo (Corces et al., 2016; Cusanovich et al., 2018; Lake et al., 2017; Preissl et al., 2018), to identify cell types and find cell type specific epigenomic signatures. Here we have shown that scATAC-seq is informative to report dynamic changes in chromatin accessibility during a time series experiment, in this case after a transcription factor perturbation. Using cisTopic we found that knockdown of SOX10 causes a fast decline of accessibility of functional SOX10 binding sites in melanoma cells, which yielded a conserved topic of around 1000 enhancers with SOX10 binding sites. Furthermore, our analysis also revealed differences in the dynamics and quantitative aspects between the cell lines. Altogether, we showed that SOX10 is a chromatin modifier and that, with the resolution of this experiment, chromatin dynamics during the EMT-like state transition occurs homogeneously across all cells of the same cell line.

We found a core SOX10 topic that is shared across a panel of melanoma cultures, as well as in melanocytes. In the melanocyte lineage, SOX10 is known as a lineage factor, together with MITF, TFAP2A, and PAX3 (Hoek et al., 2006; Scholl et al., 2001; Seberg et al., 2017; Shakhova et al., 2012). Of these transcription factors, Random Forest modelling identified the TFAP2A motif as the most informative feature, allowing to discriminate SOX10 binding in melanocytes versus other cell types, such as oligodendrocytes. In oligodendrocytes, known co-regulatory factors include OLIG1/2 (Yu et al., 2013; Zhou and Anderson, 2002). Indeed, the OLIG1/2 E-box motifs are highly informative for the classification of SOX10 binding sites in oligodendrocytes. This principle of TF cooperativity to activate enhancers in a cell-type specific manner, was confirmed by comparing these SOX10 cis-topics with a SOX9 cis-topic found in astrocytes, which share an identical SOX dimer motif with the SOX10 cis-topics. In this case, Random Forest feature selection and classification resulted in NFIA/B as the most informative cooperative motif. We were intrigued by the observation that a subset of SOX10 enhancers (17%) are shared between these cell types. SOX10 may bind strongly to these regions as SOX dimer motif scores are higher in these regions, and cofactor motifs lower. These observations lead to an enhancer model where one transcription factor has a probabilistic spectrum of binding modalities, from pioneering to cooperativity.

In conclusion, we introduce a new concept in the field of single-cell regulatory genomics, namely the cis-regulatory topic, analogous to topics in literature. We provide an easy-to-use R/Bioconductor package, called cisTopic, to discover and interpret regulatory topics and cell states from any type of single-cell epigenomics data. We believe cisTopic provides a valuable component in the analysis of large-scale single-cell epigenomics data sets, as it jointly optimises cell clustering and enhancer categorization, to identify subpopulations of cells based on shared epigenomic landscapes.

## Methods

### cisTopic workflow

cisTopic consists of 4 main steps: (1) generation of a binary accessibility matrix as input for Latent Dirichlet Allocation (LDA); (2) LDA and model selection; (3) cell state identification using the topic-cell distributions from LDA and (4) exploration of the region-topic distributions. cisTopic is available as an R/Bioconductor package at: http://github.com/aertslab/cistopic.

#### Input and binarisation

The input for cisTopic is a binary accessibility matrix, which can be built from a set of single-cell bam files and a bed file with candidate regulatory regions (e.g. from peak calling on the aggregate or the bulk profile). In the case of single-end reads, we count a fragment if its 5’ end falls within the region; in the case of paired end data, if any of its ends falls within the region. By default, we consider a region accessible if at least one read is found, leading to a binarised count matrix. In the case of single-cell methylation data, the matrix can be built from the beta values scores per region per cell, which can be also calculated if the user provides the methylation call files (i.e. tab-delimited files containing chromosome, position, number of methylated reads and total number of reads). By default, we consider a region methylated if the beta value is above 0.5. Note that regions have been blacklisted for potential artefacts prior to the analysis (http://mitra.stanford.edu/kundaje/akundaje/release/blacklists/).

#### Modelling via Latent Dirichlet Allocation

The next step in the cisTopic workflow is to use Latent Dirichlet Allocation (LDA) for the modelling of cis-regulatory topics. LDA allows to derive, from the original high-dimensional and sparse data, (1) the probability distributions over the topics for each cell in the data set (*θ*) and (2) the probability distributions over the regions for each topic (*ϕ*) (Blei et al., 2003). These distributions indicate, respectively, how important a regulatory topic is for a cell (*θ*), and how important regions are for the regulatory topic (*ϕ*). Here, we use a collapsed Gibbs sampler (Griffiths and Steyvers, 2004), in which we assign regions to a certain topic by randomly sampling from a distribution where the probability of a region being assigned to a topic is proportional to the contributions of that region to the topic and the contributions of that topic in a cell:

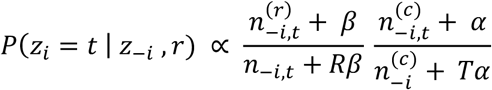

Where:

- *z_i_* is the current assignment to be made,
- *z_-i_* are the rest of assignments in the data set,
- *t* is the given topic,
- *r* is the given region,
- and *P(z_i_* = *t* | *z_-i_*, *r)* is the probability of assigning the given region *r* to a regulatory topic *t* given the rest of the assignments in the data set.
- 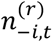 is the number of times the given region *r* is assigned to topic *t* without considering the region we want to assign,
- *β* is the Dirichlet hyperparameter of the prior distribution for the categorical distribution over regions in a topic 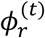. Here, we use symmetric Dirichlet priors for all topics, using 0.1 as value for *β*.
- *n*_–*i*,*t*_ is the total number of assignments to topic *t* through the data set,
- *R* is the total number of regions in the data set,
- and 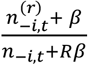 expresses the probability of region *r* under topic *t*.
- 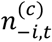 is the total number of assignments to topic *t* within the given cell *c* (without considering the region to be assigned),
- *α* is the Dirichlet hyperparameter of the prior distribution for the categorical distribution over topics in a cell *θ*^(*c*)^. Here, we use symmetric Dirichlet priors for all cells, using 50/T as value for *α*.
- 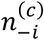 is the total number of assignments within the given cell *c*,
- *T* is the total number of topics in the model. The total number of topics has to be provided (see *Model selection)*,
- and 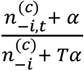 is the probability of topic *t* under cell *c*.

After enough iterations through every region in each cell in the data set, this distribution is stabilised, and assignments can be recorded. In most cases, we used 500 as burn-in and 1000 recording iterations (see *Model selection* and *Data analysis*). LDA provides two matrices, one containing the total number of assignments per topic in each cell, and another containing the total number of assignments per region to each topic. Models are built using the lda R package (Chang, 2015).

#### Model selection

For performing LDA, values for the Dirichlet priors *α* and *β*, the number of topics T and the number of iterations (burn-in and recording iterations) must be provided. We used 50/T and 0.1 for *α* and *β*, respectively, as recommended by Griffiths & Steyvers (2004). The log-likelihood per iteration in each model was plotted for confirming that the number of burn-in and recording iterations was correctly chosen (i.e. log-likelihood of the model must be stabilized when the recording of iterations starts). Several models with different number of topics were run (generally, from 5 to 50 topics; see *Data analysis*), and the optimal number of topics is selected based on the highest log-likelihood in the last iteration.

#### Cell state identification

Using the normalised topic-cell distributions (i.e. a matrix containing cells as columns, topics as rows, and normalised assignments per cell as values), cell states are visualized using dimensionality reduction methods such as tSNE (R package Rtsne (Krijthe and van der Maaten, 2017)), PCA and/or diffussion maps (R package Destiny (Angerer et al., 2016)). Hierarchical clustering with euclidean distances and ward clustering is used for the topic-cell heatmaps.

#### Topic exploration

The region-topic distributions can be explored in different ways to understand the biological nature of the regulatory topics:

##### • Enrichment of epigenomic signatures

Epigenomic signatures are intersected with the regulatory regions in the data set (by default, with at least 40% overlap) and summarized into region sets. These region sets are used, together with the normalised region-topic distributions as input for AUCell (Aibar et al., 2017). Here, we used as threshold to calculate the AUC 3% of the total number of regions in the dataset.

##### • Region annotation

Regions in the data set are annotated using the R package ChIPseeker (Yu et al., 2015). Enrichment of region types within the topics is calculated as previously explained.

##### • Topic binarisation

Representative regions of each topic are selected by rescaling the normalised region-topic assignments to the unit, and fitting a gamma distribution to these values. A threshold is given to select region above a certain probability (see *Data analysis*).

##### • Gene Ontology analysis

GO analyses was performed by using rGREAT on the binarised topics (Gu, 2018).

##### • Motif enrichment

Motif enrichment was performed using a RcisTarget (Aibar et al., 2017). cisTopic includes functions for performing motif enrichment analysis in sets of regions, rather than sets of genes. Here, we used the region-based hg19 cisTarget feather databases (v8). The cisTarget motif collection comprehends more than 20,000 PWMs obtained from JASPAR (Portales-Casamar et al., 2010), cis-bp (Weirauch et al., 2014), Hocomoco (Kulakovskiy et al., 2018), among others (Janky et al., 2014). We used a minimum fraction overlap of 0.4; a minimum Normalised Enrichment Score (NES) threshold of 3; a ROC threshold for AUC calculation of 0.005 and a threshold for visualization of 20,000. Region-based feather databases are available at: https://resources.aertslab.org/cistarget/. Motif annotation is available within the RcisTarget package.

##### • Cistrome formation

Cistromes can be formed based on RcisTarget results; by selecting the regions that pass the given thresholds. These sets of regions are linked to transcription factors based on motif annotations (direct and inferred). These cistromes are initially formed by Ctx regions (Imrichová et al., 2015), that are mapped back to the original coordinates in the data set (here, regions are mapped back if there is at least 40% of overlap).

### Validation of cisTopic

#### Simulated epigenomes from melanoma cell lines

We simulated 700 single-cell epigenomes from 14 bulk H3K27Ac ChIP-seq melanoma profiles (50 cells per bulk) by randomly sampling a given number of reads. Eleven of these bulk epigenomes were taken from Verfaille, Imrichová & Kalender-Atak et al. (2015, GSE60666); and three have been generated in this work with the same protocol and analysis pipeline. Candidate regulatory regions were defined by peak calling with MACS2 in each bulk profile (v.2.0.10, with q < 0.001 and nomodel parameters and using as control the merged control profiles of five cell lines; namely A375, MM011, MM032, MM047 and MM057) and merging of overlapping peaks. The number of reads per cell was selected randomly from the intervals corresponding to each simulation, namely 26,940-59,580 reads per cell; 8,980-19,860 reads per cell; 5,388-11,916 reads per cell and 2,694-5,958 reads per cell. For each simulation we ran cisTopic (parameters: α=50/T; β=0.1; burn-in iterations=500; recording iterations=1000) for models with a number of topics between 2 to 50 (from 2 to 30, 1 by 1; from 30 to 5, by 5). The best model in each simulation was selected based on the highest log-likelihood, resulting in selected models with 22, 22, 19 and 12 topics, from highest to lowest coverage. We binarised the topics using a probability threshold of 0.975, and performed GO enrichment analysis with rGREAT and motif enrichment analysis with RcisTarget. Latent Semantic Indexing (LSI) was performed as described by Cusanovich et al., 2015. The number of PCs selected was 7, 5, 5 and 5, for the different coverages respectively; and the first principal component was removed in all cases as it was correlated with the read depth. Values of the LSI matrix were rescaled between ±1.5. We ran chromVAR (Schep et al., 2017) with default parameters and adding the GC bias. We run scABC with default parameters, resulting in models with 14, 14, 13 and 7 landmarks (Zamanighomi et al., 2018). Rtsne was used for visualization in all cases with 50 PCs and 30 as perplexity (after testing several combinations of parameters) (Krijthe and van der Maaten, 2017). For calculating the Adjusted Rand Index, we used as ground truth the bulk epigenome of each cell and determined the cell clusters from each method using euclidean distance and ward clustering (using the cell-topic distributions matrix from cisTopic, the LSI matrix, the cistrome enrichment matrix from chromVAR and the cell-to-landmark matrix from scABC, respectively). We also tested the robustness of these methods to find rare subpopulations by reducing the number of single-cell epigenomes from 50 to 5 for 3 of these cell lines (A375, MM001 and MM099). Methods were run as previously described, and precision and recall values were calculated by using as ground truth the bulk epigenome of each cell. The cell clusters were clustered for each method using euclidean distances and ward clustering. The clusters with the highest ratio of true positives versus false positives were selected for the calculations.

#### scATAC-seq in the hematopoeitic system

We used cisTopic on a publicly available scATAC-seq data set from the hematopoeitic system (Corces et al., 2016; GSE74310), containing Leukemia Stem Cells (LSC), blasts and monocytes. We used cells with more than 784 reads per cell, resulting in a data set with 71 LSCs, 115 blasts and 77 monocytes and 296,285 regulatory regions. We ran cisTopic using α=50/T; β=0.1; burn-in iterations=200; recording iterations=1000 and models with a number of topics between 2 and 50 (by 2). The selected model had 10 topics. We binarised the topics with a probability threshold of 0.995.

#### scWGBS in the hematopoeitic system

We applied cisTopic on a publicly available scWGBS data set from the hematopoeitic system (Farlik et al., 2016; GSE87197), containing methylation calls for 18 Hematopoietic Stem Cells (HSC), 18 Multipotent Progenitors (MPP), 24 Multi-Lymphoid Progenitors (MLP), 19 Common Myeloid Progenitors (CMP) and 22 Granulocyte Macrophage Progenitors (GMP). We aggregated the methylation calls using the Ensemble regulatory regions (v78) and calculated the β values by dividing the aggregated number of methylated calls by the total number of calls, resulting in 410,037 regulatory regions. This matrix was binarised, considering as methylated regions with a β value above 0.5. We performed models using with α=50/T; β=0.1; burn-in iterations=500; recording iterations=1000; and a number of topics between 5 and 50 (by 5), resulting in a model with 10 topics to be selected. We binarised the topics with a probability threshold of 0.995 and lift-overed the regions from hg38 to hg19 before using RcisTarget.

#### scTHS-seq and scRNA-seq in the human brain

We analysed a data set from the human brain with 34,520 cells and 287,381 regulatory regions (Lake et al., 2018; GSE97942). This data set contains cells from the visual cortex, the frontal cortex and the cerebellum. We ran cisTopic with α=50/T; β=0.1; burn-in iterations=500; recording iterations=1000; and a number of topics between 5 and 50 (from 2 to 30 by 1; from 30 to 50 by 5), resulting in a model with 23 topics to be selected. We binarised the topics with a probability threshold of 0.99 and lift-overed the regions from hg38 to hg19 before using RcisTarget and rGREAT.

We filtered the scRNA-seq data from Lake et al. (2018) (GSE97930) keeping only cells with at least 800 genes expressed, resulting in a data set with 15,884 cells. SCENIC was run using default parameters (Aibar et al., 2017), resulting in a matrix with 250 regulons. Next, we mapped the regions to their closest gene, and this dictionary was used to convert the gene-based regulons to region-based regulons. These region sets were used as epigenomic signatures to determine their enrichment within the topics using AUCell as previously explained.

#### scATAC-seq during an EMT-like transition in melanoma

We generated scATAC-seq data on different time points (0, 24, 48 and 72h) for two melanoma cell lines (MM057 and MM087) upon SOX10 KD, which triggers an EMT-like cell state transition, resulting in a data set with 598 and 78,262 accessible regions (see below). We ran cisTopic with α=50/T; β=0.1; burn-in iterations=500; recording iterations=1000; and a number of topics between 5 and 50 (from 2 to 30 by 1; from 30 to 50 by 5), finding a model with 15 topics to be optimal. Topics were binarised using a probability threshold of 0.975 before RcisTarget and rGREAT analyses.

### Random forest modelling

SOX region sets were derived by merging the SOX cistromes found in the astrocytes, oligodendrocytes and shared melanoma topics, respectively. Regions were scored with Cluster Buster (Frith et al., 2003) using known and *de novo* motifs, and the value for the best CRM score in the sequence was used as feature. Known motifs were taken from the cisTarget (Herrmann et al., 2012; Imrichová et al., 2015) motif collection (see above); while *de novo* motifs were found by comparing non-overlapping regions between the SOX cistromes in a pairwise manner with Homer (Heinz et al., 2010) and RSAT *peak-motifs* (Thomas-Chollier et al., 2011, 2012). DNA shape measurements were also included as features. They were derived from models found in GBshape and Kaplan et al. (Chiu et al., 2015; Kaplan et al., 2009), using the average value between ±250 bp from the centre of the region. Comparisons were done in a pairwise manner. Per comparison, an initial selection of features was performed using a likelihood ratio test (FDR adjusted p-value < 0.05), as implemented in MAST (Finak et al., 2015). These initial features were further pruned using Boruta (Kursa and Rudnicki, 2010), using default parameters. Boruta features that represented similar motifs and showed strong correlation were merged into one Hidden Markov Model score using Cluster-Buster (Frith et al., 2003). Random forest models were performed with each set of features (namely Boruta features, merged features and a random classifier) using the randomForest R package (Liaw and Wiener, 2001). Representative rules were extracted using the package inTrees (Deng, 2014), with default parameters.

### Cell culture and treatment

The two melanoma cultures (MM057 and MM087) are short-term cultures derived from patient biopsies (Gembarska et al., 2012; Verfaillie et al., 2015). Cells were kept at 37°C, with 5% CO_2_ and were maintained in Ham’s F10 nutrient mix (Thermo Fished Scientific) supplemented with 10% fetal bovine serum (FBS; Invitrogen) and 100 μg ml^-1^ penicillin/streptomycin (Thermo Fished Scientific). SOX10 KD was performed using a SMARTpool of four siRNAs against SOX10 (SMARTpool: ON-TARGETplus SOX10 siRNA, number L017192-00-0005, Dharmacon) at a concentration of 20nM using as medium Opti-MEM (Thermo Fished Scientific) and omitting antibiotics. The cells were incubated for 24, 48 or 72 hours before processing.

### OmniATAC-seq

#### Data generation

OmniATAC-seq was performed as described previously (Corces et al., 2017). Cells were washed, trypsinised, spun down at 1000 RPM for 5 min to remove the medium and resuspended in 1 mL. Cells were counted and experiments were only continued when a viability of above 90% was observed. 50,000 cells were pelleted at 500 RCF at 4°C for 5 min, medium was carefully aspirated and the cells were washed and lysed using 50 uL of cold ATAC-Resupension Buffer (RSB) (see Corces et al., 2017 for composition) containing 0.1% NP40, 0.1% Tween-20 and 0.01% digitonin by pipetting up and down three times and incubating the cells for 3 min on ice. The lysis was washed out by adding 1 mL of cold ATAC-RSB containing 0.1% Tween-20 and inverting the tube three times. Nuclei were pelleted at 500 RCF for 10 min at 4°C, the supernatant was carefully removed and nuclei were resuspended in 50 uL of transposition mixture (25 uL 2x TD buffer (see Corces et al., 2017 for composition), 2.5 uL transposase (100 nM), 16.5 uL DPBS, 0.5 uL 1% digitonin, 0.5 uL 10% Tween-20, 5 uL H2O) by pipetting six times up and down, followed by 30 minutes incubation at 37°C at 1000 RPM mixing rate. After MinElute clean-up and elution in 21 uL elution buffer, the transposed fragments were pre-amplified with Nextera primers by mixing 20 uL of transposed sample, 2.5 uL of both forward and reverse primers (25 uM) and 25 uL of 2x NEBNext Master Mix (program: 72°C for 5 min, 98°C for 30 sec and 5 cycles of [98°C for 10 sec, 63 °C for 30 sec, 72°C for 1 min] and hold at 4°C). To determine the required number of additional PCR cycles, a qPCR was performed (see Buenrostro et al., 2015 for the determination of the number of cycles to be added). The final amplification was done with the additional number of cycles, samples were cleaned-up by MinElute and libraries were prepped using the KAPA Library Qunatificaton Kit as previously described (Corces et al., 2017). Samples were sequenced on a NextSeq500 High Output chip, generating between 41 and 70 million reads per sample.

#### Data processing

Adapter sequences were trimmed from the fastq files using fastq-mcf (as part of ea utils; v1.04.807). Read quality was then checked using FastQC (v0.11.5). Reads were mapped to the human genome (hg19-Gencode v18) using STAR (v2.5.1) applying the parameters --alignIntronMax 1 and --aslignIntronMin 2. Mapped reads were filtered for quality using SAMtools (v1.2) view with parameter –q4, sorted with SAMtools sort and indexed using SAMtools index. Peaks were called using MACS2 (v2.1.1) callpeak using the parameters --nomodel and --call-summits on the 8 conditions separately. A count matrix was generated by using featureCounts (as part of Subread; v1.4.6) of all separate bam files on the merged peak file (after conversion of the merged peak bed file to a gff format using a custom script). Normalised bedGraphs were produced by genomeCoverageBed (as part of bedtools; v2.23.0) using as scaling parameter (-scale) size factors obtained from DEseq2 (v1.18.1). BedGraphs were converted to bigWigs by the bedtools suit functions bedSort to sort the bedGraphs, followed by bedGraphToBigWig to create the bigWigs, which were used in IGV for visualisation.

### scATAC-seq

#### Data generation

scATAC-seq was performed using the Fluidigm C1 system as described before (Buenrostro et al., 2015). Briefly, cells were trysinised, spun down (1000 RPM, 5 min), medium was removed and cells were resuspended in fresh medium and passed through a 40 um filter, counted and diluted till 200,000 cells per mL. Cells were loaded (using a 40:60 ratio of RGT:cells) on a primed Open App IFC (10-17 um, the protocol for ATAC-seq from the C1 Script Hub was used). After cell loading, the plate was visually checked under a microscope and the number of cells in each of the capture chambers was noted. Next, the “Sample prep” was performed on the Fluidigm C1 during which the cells underwent lysis and ATAC-seq fragments were prepared. In a 96-well plate, the harvested libraries were amplified in a 25 uL PCR reaction. The PCR products were pooled and purified on a single MinElute PCR purification column for a final library volume of 15 uL. Quality checks were performed using the Bioanalyzer high sensitivity chips. Fragments under 150 bp were removed by bead-cleanup using AMPure XP beads (1.2x bead ratio) (Beckman Coulter). All scATAC-seq libraries were sequenced on a HiSeq4000 paired-end run, generating a median of 170,769 raw reads per single cell.

#### Data processing

The reads from scATAC-seq samples were first cleaned for adapters using fastq-mcf using fastq-mcf (as part of ea utils; v1.1.2-686). Read quality was then checked using FastQC (v0.11.5). Paired-end reads were mapped to the human genome (hg19-Gencode v18) using STAR (v2.5.1) applying the parameters --alignIntronMax 1, --aslignIntronMin 2 and --alignMatesGapMax 2000. Mapped reads were filtered for quality using SAMtools (v1.2) view with parameter –q4, sorted with SAMtools sort and indexed using SAMtools index. Duplicates were removed using Picard (v1.134) MarkDuplicates using OPTICAL_DUPLICATE_PIXEL_DISTANCE=2500. To filter out cell of bad quality, transcription start site aggregation plots were made using a custom script and cell having a low signal-to-noise profile were removed from further analyses. This lead to a final of 598 good quality cells over 8 Fluidigm C1 runs. Bam files of good quality single cells were aggregated per condition and peaks were called on these aggregated samples using MACS2 (v2.1.1) callpeak using the parameters --nomodel and --call-summits. The peak files per condition were merged (78,661 peaks in total before blacklisting) and blacklisted using the blacklisted regions of hg19 listed on http://mitra.stanford.edu/kundaje/akundaje/release/blacklists/hg19-human/ (Anshul Kundaje), leading to a total of 78,262 peaks after blacklisting. This peak file was used, together with the bam files of the good single cells as, input for cisTopic. To visualise the aggregated cells per sample, normalised bedGraphs were produced by genomeCoverageBed (as part of bedtools; v2.23.0) using as scaling parameter (-scale) size factors obtained from DEseq2 (v1.18.1). BedGraphs were converted to bigWigs by the bedtools suit functions bedSort to sort the bedGraphs, followed by bedGraphToBigWig to create the bigWigs.

### Luciferase assays

The DCT (chr13:95131958-95132420) and EDNRB (chr13:78427800-78428233) regulatory regions were defined based on the peaks obtained in our scATAC-seq experiment. The regions were scored with Cluster-Buster (Frith et al., 2003) for the SOX dimer motif (transfac_pro＿ MM08838) and the identified motifs were disrupted by two point mutations (AC<u>A</u>aagnnncctt<u>T</u> to AC<u>C</u>aagnnncctt<u>G</u>), manually changing these nulceotides in the fasta sequences. Similarly, the wild-type regions were scored for Eboxes (most probably linked to MITF) and these were disrupted by two point mutations (<u>C</u>ANNT<u>G</u> to TANNT<u>A</u>), taking care that no SOX motifs were consequently disrupted. Lastly, we modified the inner two nucleotides of the Eboxes from the putative MITF Ebox (CA<u>CG</u>TG) to the putative Olig Ebox (CA<u>GC</u>TG). The wild-type sequence and the mutated sequences were synthetically generated, together with specific cloning sites, via gBlocks (IDT). The fragments were cloned into a pGL4.23[luc2/minP] vector (Promega) using cohesive-end restriction cloning. Clones were checked by Sanger sequencing for the correct mutation. Luciferase assays were performed three times in triplicate for each plasmid. Cells seeded at ~80% confluency were transfected with 400 ng of the luciferase reporter plasmid and 40 ng of Renilla plasmid (Promega) using lipofectamine 2000 (Invitrogen). Luciferase activity of each variant was measured using the Dual-Luciferase Reporter Assay (Promega) and was normalised against the Renilla luciferase activity. We performed a two-sided t-test with unequal variance and calculated the standard deviation.

### Publicly available data used in this work

Raw fastq files of DNAseI-seq on penis foreskin melanocytes primary cells were downloaded from NCBI’s Gene Expression Omnibus (Edgar et al., 2002) through GEO accession number GSE18927 (GSM774243) and was mapped on the human genome (hg19-Gencode v18) using STAR (v2.5.1). SOX10 ChIP-seq and MITF ChIP-seq were downloaded as raw fastq files from GEO GSE61965 and were mapped to the human genome using Bowtie2 (v2.1.0) and peaks were called by MACS2 (v2.1.1). TFAP2 ChIP-seq data in human primary melanocytes was retrieved from Seberg et al., 2017 (GSE67555). FAIRE-seq, H3K27Ac-seq and RNA-seq data on the melanoma lines (GSE60666) were processed as mentioned in Verfaillie et al., 2015.

For the simulations of single cells from bulk melanoma cell line epigenomes, we used the H3K27Ac data from Verfaillie et al., 2015 (GEO GSE60666). scATAC-seq data from the hematopoeitic system (Corces et al., 2016), was retrieved from GEO GSE74310; scWGBS data in the hematopoeitic system (Farlik et al., 2016) was obtained from GEO GSE87197; and scTHS-seq and scRNA-seq data from the human brain (Lake et al., 2017) was downloaded from GEO GSE97942 and GEO GSE97930, respectively.

## Data availability

The data generated for this study have been deposited in NCBI’s Gene Expression Omnibus and are accessible through GEO Series accession number GSE114557.

## Code availability

cisTopic is available as an R package at: http://github.com/aertslab/cistopic.

## Acknowledgements

This work is funded by an ERC Consolidator Grant to S. Aerts (724226_cis-CONTROL); by the Special Research Fund (BOF) KU Leuven (grants PF/10/016 to S. Aerts), the Harry J. Lloyd Charitable Trust, the Foundation Against Cancer (2016-070; to S. Aerts), PhD fellowships from the F.W.O. (L.M.) and a postdoctoral research fellowship from Kom op tegen Kanker (Stand up to Cancer), the Flemish Cancer Society (J.W.). Computing was performed at the Vlaams Supercomputer Center (VSC). The funders had no role in study design, data collection and analysis, decision to publish or preparation of the manuscript. The authors thank Jean-Christophe Marine, Florian Rambow, and Michael Dewaele for helpful discussions; and Blue Lake and Kun Zhang for the information provided regards the human brain data. The authors also thank various groups that make curated position weight matrices publicly available, including T. Hughes (cis-bp), M. Bulyk (Uniprobe), A. Mathelier (Jaspar), V. Makeev (Hocomoco) and many others.

## Supplementary Figures

**Fig S1:** cisTopic on simulated single-cell epigenomes from 14 bulk H3K27Ac profiles from different melanoma cell lines using medium-high coverage (8,980-19,860 reads per cell).

**Fig S2:** cisTopic detects rare subpopulations with higher accuracy and precision than other methods, namely LSI and chromVAR.

**Fig S3:** cisTopic reveals differentiation dynamics in the hematopoietic system and oncogenesis.

**Fig S4:** cisTopic reconstructs a differentiation hierarchy in the hematopoeitic system from scWGBS data.

**Fig S5:** cisTopic model selection in the human brain data set.

**Fig S6:** cisTopic on the human brain.

**Fig S7:** Summarised cisTopic results for Lake et al. (2017).

**Fig S8:** Accessibility profiles of the cerebellum (CBL), visual cortex (BA17) and frontal cortex (BA9) in the vicinity of *NEUROD1*.

**Fig S9:** Enrichment of SCENIC regulons within topics in the human brain.

**Fig S10:** Correlation between single-cell and bulk ATAC-seq.

**Fig S11:** Loss of accessibility during the EMT-like transition at known SOX10-bound and -activated melanocyte-like regulatory regions and gain of accessibility at mesenchymal-like regions.

**Fig S12:** cisTopic identifies 15 regulatory topics involved in melanoma phenotype switching.

**Fig S13:** Topics identified by cisTopic on melanoma scATAC-seq data.

**Fig S14:** Chromatin accessibility dynamics per regulatory topic.

**Fig S15:** Validation of melanocyte-like and invasive topics using ChIP-seq.

**Fig S16:** General and cell-line specific SOX10 regulatory topics govern the melanocyte-like state.

**Fig S17:** Melanoma SOX10 enhancers are accessible in melanocytes.

**Fig S18:** DNA shape features profiles for the astrocyte, oligodendrocyte and melanoma SOXE cistromes.

**Fig S19:** TFAP2A and MITF ChIP-seq peaks overlap specifically with melanoma SOX cistrome regions.

**Fig S20:** AP-1 and TFAP2 members are uniquely expressed in melanoma as compared to oligodendrocytes and astrocytes.

**Fig S21:** Correlation heatmap between the DNA shape features measurements of the SOXE cistromes.

**Fig S22:** Comparison of astrocyte specific SOXE regions with melanoma and oligodendrocytes SOXE specific regions, respectively.

**Fig S23:** Mutating MITF motifs in a melanoma-specific SOX10 enhancer destroys its activity.

## Supplementary Tables

**Table S1:** Comparison of current experimental protocols for performing single-cell epigenomics assays.

**Table S2:** Comparison between current bioinformatics methods for analysing single-cell ATAC-seq data.

